# Oxycodone self-administration activates the mitogen-activated protein kinase/mitogen- and stress-activated protein kinase (MAPK-MSK) signaling pathway in the rat dorsal striatum

**DOI:** 10.1101/2020.08.31.276253

**Authors:** Christopher A. Blackwood, Michael T. McCoy, Bruce Ladenheim, Jean Lud Cadet

## Abstract

To identify signaling pathways activated by oxycodone self-administration (SA), Sprague-Dawley rats self-administered oxycodone for 20 days using short-access (ShA, 3 h) and long-access (LgA, 9 h) paradigms. Animals were euthanized two hours after SA cessation and dorsal striata were used in post-mortem molecular analyses. LgA rats escalated their oxycodone intake and separated into lower (LgA-L) or higher (LgA-H) oxycodone takers. LgA-H rats showed increased striatal protein phosphorylation of ERK1/2 and MSK1/2. Histone H3, phosphorylated at serine 10 and acetylated at lysine 14 (H3S10pK14Ac), a MSK1/2 target, showed increased abundance only in LgA-H rats. RT-qPCR analyses revealed increased AMPA receptor subunits, *GluA2* and *GluA3* mRNAs in the LgA-H rats. *GluA3*, but not *GluA2*, expression correlated positively with changes in pMSK1/2 and H3S10pK14Ac. Our findings indicate that escalated oxycodone SA results in MSK1/2-dependent histone phosphorylation, which promoted increases in striatal gene expression. Our observations offer novel avenues for pharmacological interventions against oxycodone addiction.

## Introduction

The opioid epidemic remains a public health crisis ^1,2^. This is related, in part, to the over-prescription of the opioid agonist, oxycodone, for pain management ^3–6^. Its illicit abuse has also contributed to the high number of overdose-related deaths ^7,8^. Other complications of oxycodone use disorder include moderate to severe withdrawal symptoms ^1^ and repeated episodes of relapses during attempts to quit through psychological or pharmacological interventions ^9^. Chronic use of opioid drugs is also accompanied by cognitive deficits ^10^ and post-mortem evidence of neuropathological abnormalities in the brain ^11^. These biopsychosocial complications make the development of effective treatment of paramount importance.

Pharmacological approaches to treat opioid use disorders (OUDs) have mainly included the use of agents that interact with opioids receptors ^3,12,13^. Upon activation, opioid receptors transmit signals to the nucleus via intracellular events that involve modulation of some kinase cascades ^14–16^, with consequent changes in gene expression ^17,18^. Because of several of these experiments had done using *in vitro* systems, it was important to address the potential neurobiological impact of repeated oxycodone self-administration in humans by using model systems that better mimic human conditions. Therefore, we decided to use the model of escalated oxycodone self-administration (SA) in rats ^19,20^ to identify potential biochemical and molecular pathways that might be perturbed by this drug.

Herein, we used that model to measure alterations in various proteins that may impact the flow of intracellular signals from the mu opioid receptor consequent to its repeated interactions with oxycodone during a drug SA experiment. The rat dorsal striatum was dissected and processed for biochemical and molecular analyses because this structure is thought to play essential roles in the manifestation of habitual drug taking behaviors ^21–23^. Thus, we report that the mitogen-activated protein kinase (MAPK)/mitogen- and stress-activated protein kinase (MSK) signaling cascade is activated preferentially in rats that consume large quantities of oxycodone over a period of 20 days. This was manifested by increased phosphorylation of extracellular signal-regulated kinases 1/2 (ERK1/2), MSK1/2, and increased abundance of histone H3 phosphorylated at serine 10 and acetylated at lysine 14 (H3S10pK14Ac). Altogether, these findings implicate the role of MAPK/MSK pathway and histone H3 phosphoacetylation in opioid use disorder.

## Materials and Methods

### Intravenous surgery

We used male Sprague Dawley rats (Charles River, Raleigh, NC, USA), weighing 350-400 g before surgery and housed on a 12 h reversed light/dark cycle with food and water freely available. All procedures followed the guidelines outlined in the National Institutes of Health (NIH) Guide for the Care and Use of Laboratory Animals (eighth edition, https://guide-for-the-care-and-use-of-laboratory-animals.pdf) as approved by the NIDA (National Institute of Drug Abuse) Animal Care and Use Committee at the Intramural Research Program (IRP). Catheter implantations were performed as previously described ^24^. Briefly, we anesthetized the rats with ketamine (50 mg/kg) and xylazine (5 mg/kg). Polyurethane catheters (SAI Infusion Technologies, Lake Villa, IL) were inserted into the jugular vein. The other end of the catheter was attached to a modified 22-gauge cannula (Plastics One, Roanoke, VA) that was mounted to the back of each rat. The modified cannulas, which served as infusion ports for the catheters, were connected to a fluid swivel (Instech, Plymouth, PA) via polyethylene-50 tubing that was protected by a metal spring. When the infusion ports were not used they were sealed using dust caps (PlasticOne, Roanoke, VA). Thereafter, the catheters were flushed every 48 h with gentamicin (0.05 mg/kg, Henry Schein, Melville, NY) in sterile saline to maintain patency. Intraperitoneal injection of buprenorphine (0.1 mg/kg) was used post-surgery to relieve pain.

### Apparatus

Rats were trained in Med Associates SA chambers located inside sound-attenuated cabinets and controlled by a Med Associates System (Med Associates, St Albans, VT) as previously described ^19^. In brief, each chamber was equipped with two levers located 8.5 cm above the grid floor. Presses on the retractable active lever activated the infusion pump and tone-light cue. Presses on the inactive lever had no reinforced consequences.

### Training phase

Rats (n=42) were randomly assigned to either saline (Sal) (n=8) or oxycodone (n=33) conditions. Rats were trained to self-administer oxycodone-HCL (NIDA Pharmacy, Baltimore, MD) for one 3 h daily session for the short-access (ShA) condition (n=10) or one to three 3 h sessions for long-access (LgA) condition (n=23) (Fig. 1A). For the LgA group, the 3 h sessions were separated by 30 mins intervals from day 6 to day 20 (Fig. 1A). Lever presses were reinforced using a fixed ratio-1 with a 20-s timeout accompanied by a 5-s compound tone-light cue. We used a scheduling pattern of 5 days of drug SA and 2 days off to control for weight loss, a common side effect of oxycodone intake in laboratory animals ^25^. Rats self-administered oxycodone at a dose of 0.1mg/kg per infusion over 3.5-s (0.1 ml per infusion). The house light was turned off, and the active lever retracted at the end of the 3 h session.

**Figure 1.**
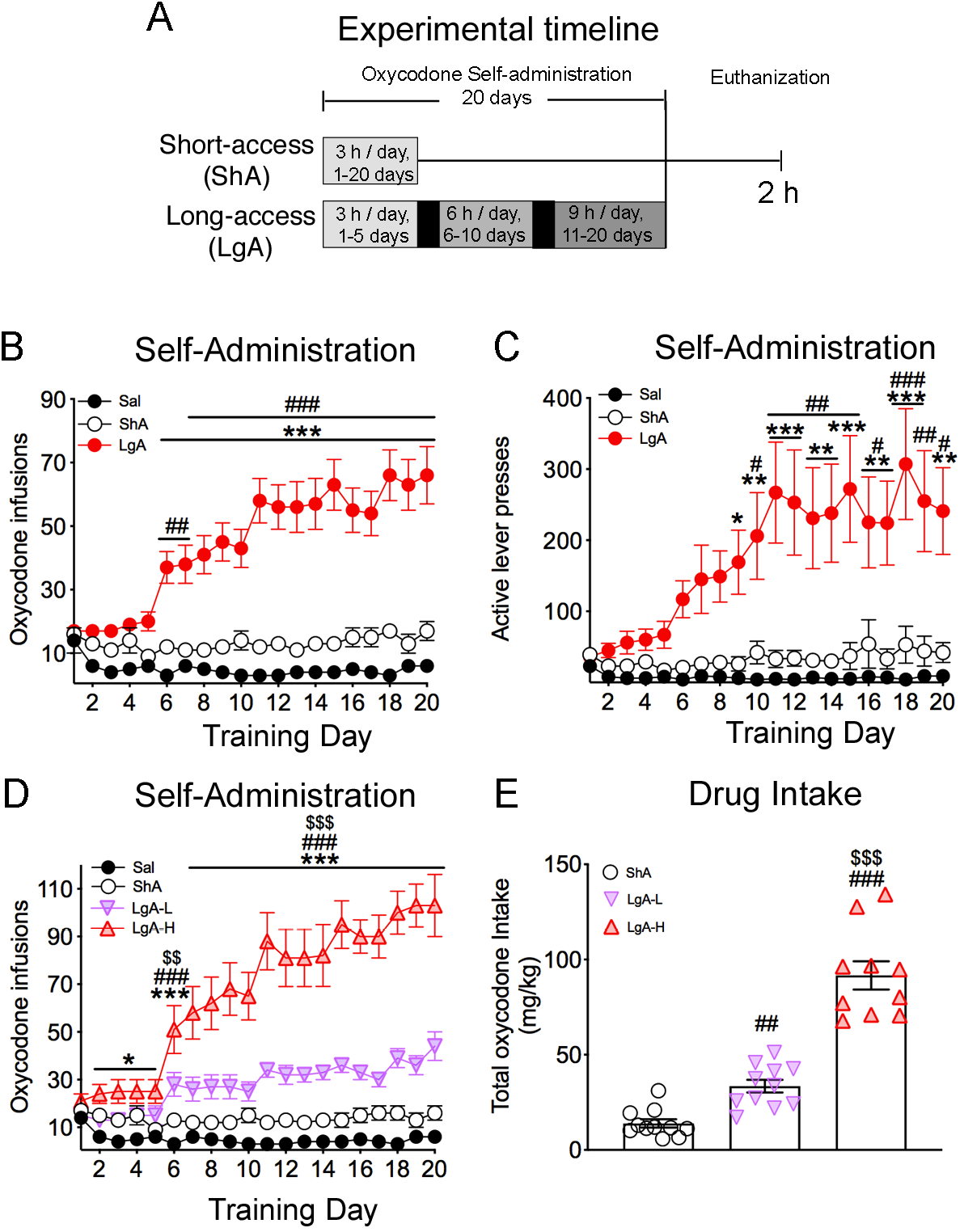
Rats exposed to long-access, but not short-access, oxycodone SA escalate their drug intake. **(A)** Experimental timeline of oxycodone self-administration (SA) training. Rats self-administered oxycodone using either short-access (ShA) (n=10) (trained for 3 h for 20 days) or long-access (LgA) (n=21-23) (trained for 3 h for 1-5 days, 6 h for 6-10 days, then 9 h for 11-20 days) paradigms. **(B)** LgA rats escalate their intake of oxycodone after the first 5 days of SA training. **(C)** LgA rats show significant increases in active lever presses during SA training. **(D)** LgA-H rats show two distinct intake phenotypes, high (LgA-H) (n=10) and low (LgA-L) (n=13) oxycodone takers, during the escalation phase. **(E)** LgA-H rats took substantially more oxycodone than LgA-L and ShA rats. Key to statistics: *, **, *** = p < 0.05, 0.01, 0.001, respectively, in comparison to Sal rats; #, ##, ### = p < 0.05, 0.01, 0.001, respectively, in comparison to SHA rats; $$, $$$ = p < 0.01, 0.001 in comparison to LgA-L rats. Stats were performed by either one-way or two-way ANOVA followed by Bonferroni or Fisher’s PLSD post hoc test.

### Tissue Collection

Rats were euthanized 2 h after training day 20. Dorsal striata tissue was dissected as previously described ^26^. In brief, we used stereotaxic coordinates (A/P +2 to −2 mm bregma, M/L ±2 to 5 mm, D/V −3 to −6 mm) according to the rat atlas ^27^ and we used the position of anatomical structures (corpus callosum and lateral ventricles) for further accuracy. In brief, the dorsal striata was removed from the skulls and snap frozen on dry ice. Tissue was later used for western blotting and quantitative RT-PCR experiments.

### Western blotting

Western blotting was conducted as previously described ^19^. Ten – twenty μg of lysate was prepared in solutions that contained 1x NuPage LDS Sample Buffer (ThermoFisher Scientific, Waltham, MA), and 1% β-Mercaptoethanol. Protein samples were heated to 70°C and loaded on 3-8% Tris-Acetate Protein Gels (ThermoFisher Scientific, Waltham, MA) or NuPAGE 4-12% Bis-Tris Protein Gels (ThermoFisher Scientific, Waltham, MA). Proteins were electrophoretically transferred on the Trans-Blot® Turbo™ system (Bio-Rad, Hercules, CA). Membrane blocking, antibody incubations, and chemiluminescence reactions were performed according to the manufacturer’s instructions. Primary and secondary antibodies are listed in Supplemental Table S1. Supplemental Table S1 also includes Research Resource Identifiers (RRIDs) where antibodies were previously validated. All antibodies ran at approximate predicted sizes according to manufacturer’s instructions. Cyclophilin B or alpha-tubulin was used as loading controls. Following secondary antibody incubation, ECL clarity (Bio-Rad, Hercules, CA) was used to visualize gel bands on ChemiDoc Touch Imaging System (Bio-Rad, Hercules, CA), and intensities were quantified with Image Lab version 6.0 (Bio-Rad, Hercules, CA) software.

### Quantitative PCR

Total RNA was collected as previously described ^19^. PCR experiments were performed using the LightCycler 480 II (Roche Diagnostics, Indianapolis, IN) with iQ SYBR Green Supermix (Bio-Rad, Hercules, CA). Primers were purchased from Johns Hopkins University (Baltimore, MD) Synthesis and Sequence Facility. Primer sequences are listed in Supplementary Table S2. The data was normalized to *Qaz1* or *B2m* reference genes. The standard curve method was used to analyze data and the results are reported as fold change relative to Sal.

### Statistical Analyses

Behavioral data were analyzed using either one-way or two-way analysis of variance (ANOVA) as previously described ^19^. In brief, dependent variables were the number of oxycodone infusions on training days. Independent variables were between-subject factor reward types (Sal, ShA, LgA-L, LgA-H), within-subject factor SA day (training days 1-20), and their interactions. If the main effects were significant (p < 0.05), Bonferroni post hoc tests were used to compare reward types on each training day. Biochemical data were analyzed using one-way ANOVA followed by the Fisher’s PLSD post hoc test. Regression analyses were performed using the correlation function in Prism version 8.3.0 (GraphPad Software, San Diego, CA). Statistical significance for all hypothesis tests was set at p < 0.05. Behavioral and biochemical data were analyzed with Prism version 8.3.0 (GraphPad Software, San Diego, CA).

## Results

### Long-access self-administration leads to escalated oxycodone intake in rats

Figure 1 shows the experimental timeline and behavioral results for oxycodone SA. As described in details under methods, rats were given short-access (ShA) or long-access (LgA) to oxycodone during the experiment ^19^. The repeated-measures ANOVA for reward earned included the between-subject factor, groups (Saline, ShA, LgA), the within-subject factor of SA days (training days 1-20), and the group × day interaction. This analysis showed statistically significant effects of group [F_(2, 890)_ = 307.5, p < 0.001], day [F_(19, 890)_ = 3.016, p < 0.0001], and significant group × day interaction [F_(38, 890)_ = 3.958, p < 0.0001]. A comparison of the LgA rats to Saline rats showed that the LgA rats increased their oxycodone intake substantially after training day 5 compared to Saline rats [F_(1, 507)_ = 35, p < 0.0001; Fig. 1B], with there being significant increases in the number of active lever presses [F_(1, 507)_ = 87, p < 0.0001; Fig. 1C] by LgA rats during the drug SA experiments. As previously reported ^19^, LgA rats could be further divided into two SA phenotypes depending on whether they took high (LgA-H) and lower (LgA-L) amounts of oxycodone (Fig. 1D). Figure 1E shows that the LgA-H rats consumed significantly more oxycodone than the ShA and LgA-L rats [F_(2, 29)_ = 85.00, p < 0.0001].

### Effects of early withdrawal and oxycodone SA on the activation of PKC

Rats were euthanized 2 hours after cessation of oxycodone SA and their dorsal striata were used in Western Blot analyses of several phospho-proteins involved in the MAPK/MSK signaling pathway. The kinase, PKC, is known to be involved in the MAPK signaling cascade stimulated by opioid receptors ^28–30^. Supplementary Figure S1 shows the effects of oxycodone SA on PKC and pPKC protein expression in the dorsal striata of rats euthanized at 2 h after cessation of drug SA. There were no significant changes in striatal PKC protein levels [F_(3, 18)_ = 0.73, p = 0.540; Supplementary Fig. S1]. However, changes in phosphorylated PKC (pPKC) abundance trended towards significance [F_(3, 18)_ = 2.82, p = 0.068] in LgA-H rats, with planned tests showing small increases in LgA-H in comparison to Saline and ShA rats (Supplementary Fig. S1). Importantly, pPKC/PKC ratios were significantly increased [F_(3, 18)_ = 4.45, p = 0.0165] in the LgA-H group compared to the other groups (Supplementary Fig. S1). Regression analysis revealed a significant positive correlation between pPKC/PKC ratios and the amount of total oxycodone taken during the SA experiment (Supplementary Fig. S1). By contrast, there were no changes in protein kinase A (PKA) protein expression [F_(3, 20)_ = 1.98, p = 0.1498] or pPKA abundance [F_(3, 20)_ = 2.22, p = 0.1178] (Supplementary Fig. S2), suggesting that changes in the PKA signaling pathway were not involved in oxycodone SA after 20 days of drug exposure in a manner consistent with a previous report that cAMP/PKA cascade might not be involved in the rewarding properties of morphine ^31^.

### Withdrawal from oxycodone SA increases ERK phosphorylation in LgA rats

ERK1/2 are members of MAPK kinases that are regulated by opioid drugs ^32^ and are also activated by PKC ^33^. We thus measured their expression in oxycodone-exposed rats and found no significant changes [F_(3, 20)_ = 2.18, p = 0.1224] in striatal ERK1/2 protein expression (Figs. 2A, 2B). However, there were increases [F_(3, 20)_ = 6.41, p = 0.0032] in the abundance of pERK1/2 in the LgA-H group in comparison to Saline and ShA groups (Figs. 2A, 2C). pERK/ERK ratios were also significantly increased [F_(3, 20)_ = 7.10, p = 0.0020] in the LgA-H group in comparison to the other 3 groups (Fig. 2D). Regression analysis showed oxycodone amount-dependent increases in pERK/ERK ratios (Fig. 2E).

**Figure 2.**
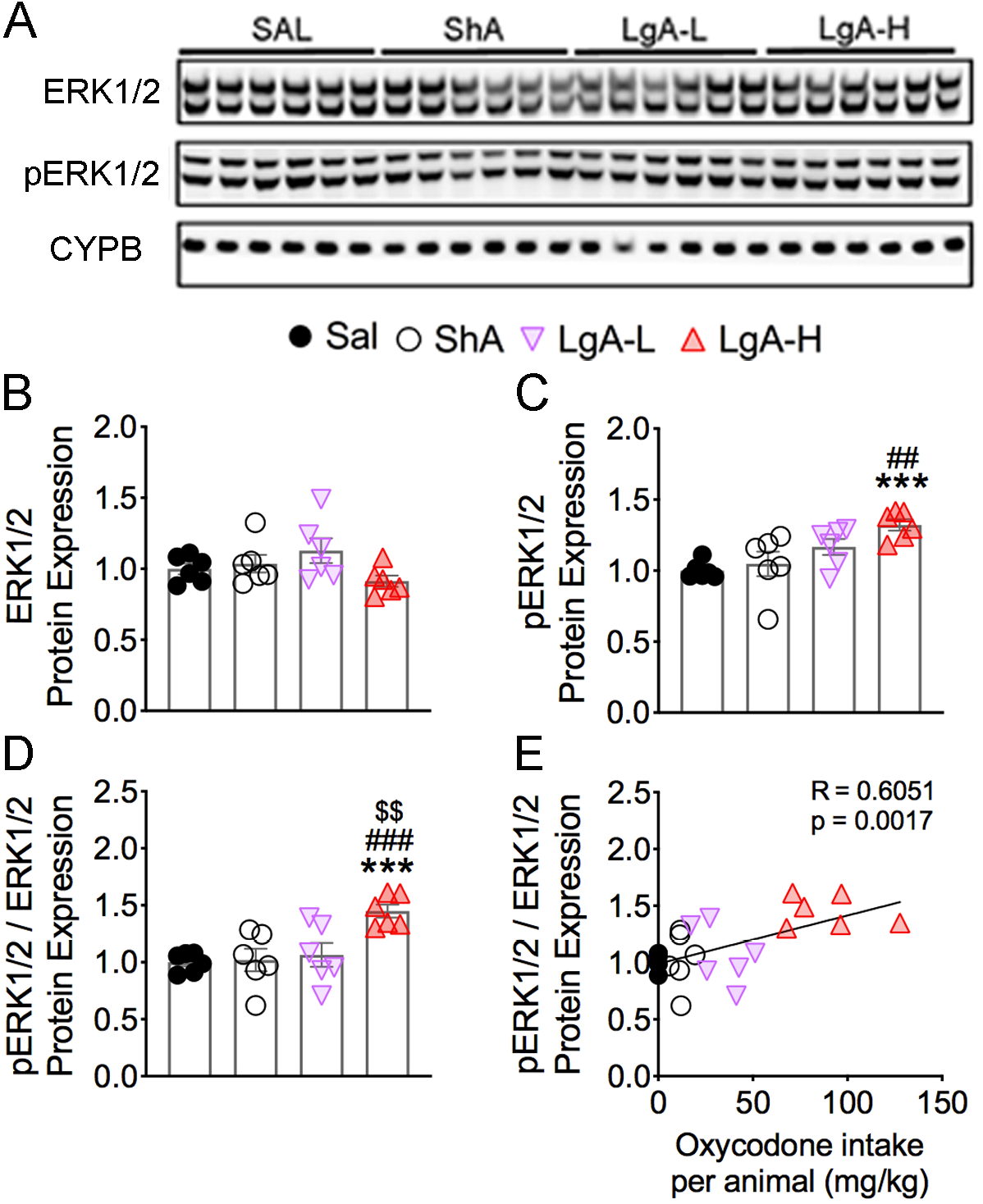
Effects of oxycodone SA on ERK1/2 phosphorylation. **(A)** Images of western blot and **(B, C)** quantification of ERK1/2, and pERK1/2. **(B)** ERK1/2 protein levels were not significantly impacted by oxycodone. **(C)** pERK1/2 abundance was increased in only LgA-H rats. **(D)** pERK1/2/ERK1/2 ratios are substantially increased in the LgA-H rats. **(E)** pERK1/2/ERK1/2 ratios correlate with amount of oxycodone taken (n=6 Sal; n=6 ShA; n=6 LgA-L; n=6 LgA-H). Key to statistics: *** = p < 0.001 in comparison to Sal rats; ##, ### = p < 0.01, 0.001, respectively, in comparison to ShA rats; $$ = p < 0.01, in comparison to LgA-L rats. Statistical analyses are as described in Figure 2.

### Effects of oxycodone and early withdrawal on MSK1 and MSK2 proteins

ERK1/2 kinases phosphorylate MSK1 and MSK2 proteins in the MAPK/MSK cascade ^34,35^. MSK1 is also activated in neurons in response to stress and neurotrophins ^36^. We therefore tested the possibility that MSK1 and MSK2 phosphorylation might be affected in oxycodone SA animals. Figure 3 shows the effects of oxycodone SA on MSK1, pMSK1, MSK2, and pMSK2 levels. MSK1 protein expression was significantly decreased [F_(3, 20)_ = 6.18, p = 0.0038] in both LgA-L and LgA-H rats in comparison to the Saline group (Figs. 3A, 3B). However, pMSK1 abundance was substantially increased [F_(3, 20)_ = 4.53, p = 0.0140] in the LgA-H group in comparison to Sal and ShA groups (Figs. 3A, 3C). In addition, pMSK1/MSK ratios were increased [F_(3, 20)_ = 16.3, p < 0.0001] in both LgA-L and LgA-H rats in comparison to Sal and ShA animals (Fig. 3D). Regression analysis revealed significant correlation between pMSK1/MSK ratios and amount of oxycodone self-administered (Fig. 3E).

**Figure 3.**
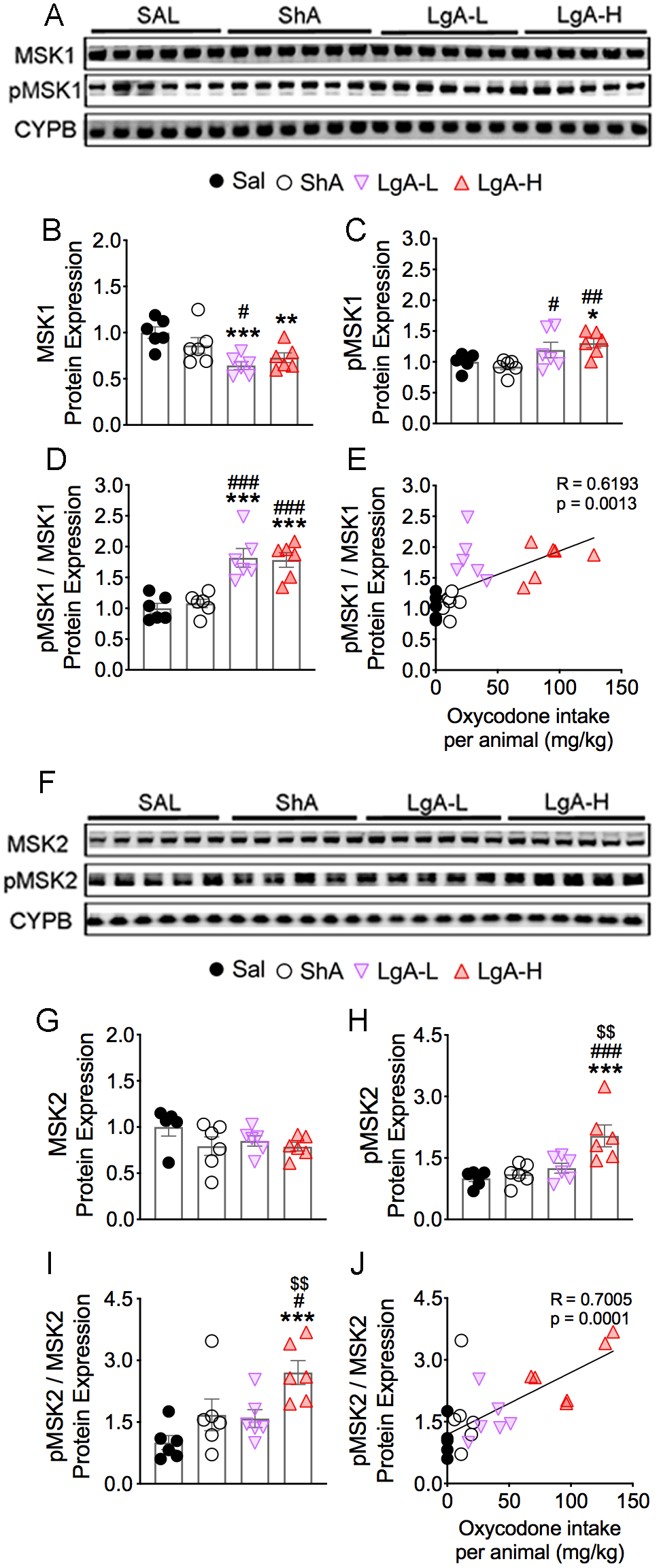
Increased MSK1 and MSK2 protein phosphorylation in LgA-H rats. **(A, F)** Images of western blot and **(B, C, G, H)** quantification of MSK1, pMSK1, MSK2, pMSK2. **(B)** MSK1 protein levels are decreased in LgA-L and LgA-H rats. **(C)** pMSK1 protein abundance is upregulated in LgA-L and LgA-H rats. **(D)** pMSK1/MSK1 ratios are increased in LgA rats; **(E)** Changes in pMSK1/MSK1 ratios are dependent on oxycodone intake. **(G)** MSK2 protein levels were not significantly affected by oxycodone. **(H**) pMSK2 abundance is increased in LgA-H rats. **(I)** pMSK2/MSK2 ratios are increased in LgA-H rats and **(J)** correlated with amount of oxycodone (n=5-6 Sal; n=6 ShA; n=6 LgA-L; n=6 LgA-H). Key to statistics: *, **, *** = p < 0.05, 0.01, 0.001, respectively, in comparison to Sal rats; #, ##, ### = p < 0.05, 0.01, 0.001, respectively, in comparison to SHA rats; $$ = p < 0.01 in comparison to LgA-L rats. Statistical analyses are as described in Fig. 2.

There were no significant changes [F_(3, 19)_ = 1.52, p = 0.2413] in MSK2 protein expression (Figs. 3F, 3G). There were, however, significant increases [F_(3, 20)_ = 8.73, p = 0.0007] in pMSK2 abundance only in LgA-H rats in comparison to other groups (Figs. 3F, 3H). pMSK2/MSK2 ratios were also significantly increased [F_(3, 20)_ = 6.50, p = 0.0030] in only LgA-H rats (Fig. 3I), with significant positive correlation observed between these ratios and amount of oxycodone taken during the SA experiment (Fig. 3J).

### Increased pCREB in LgA-H rats

Because CREB phosphorylation can be mediated by several upstream kinases that include PKC, ERK1/2 and MSKs ^37^, we examined the possibility that activation of these kinases might have led to increased pCREB after oxycodone SA. There were no significant changes [F(_3, 18)_ = 2.97, p = 0.0592] in CREB protein expression (Figs. 4A, 4B). However, the abundance of pCREB was significantly increased [F(_3, 20)_ = 5.71, p = 0.0054] in LgA-L and LgA-H groups in comparison to the Saline group (Figs. 4A, 4C). Furthermore, pCREB/CREB ratios were also substantially increased [F_(3, 20)_ = 3.93, p = 0.0235] in the LgA-H group compared to other groups (Fig. 4D), with there being a significant positive correlation between these ratios and amount of oxycodone taken (Fig. 4E).

**Figure 4.**
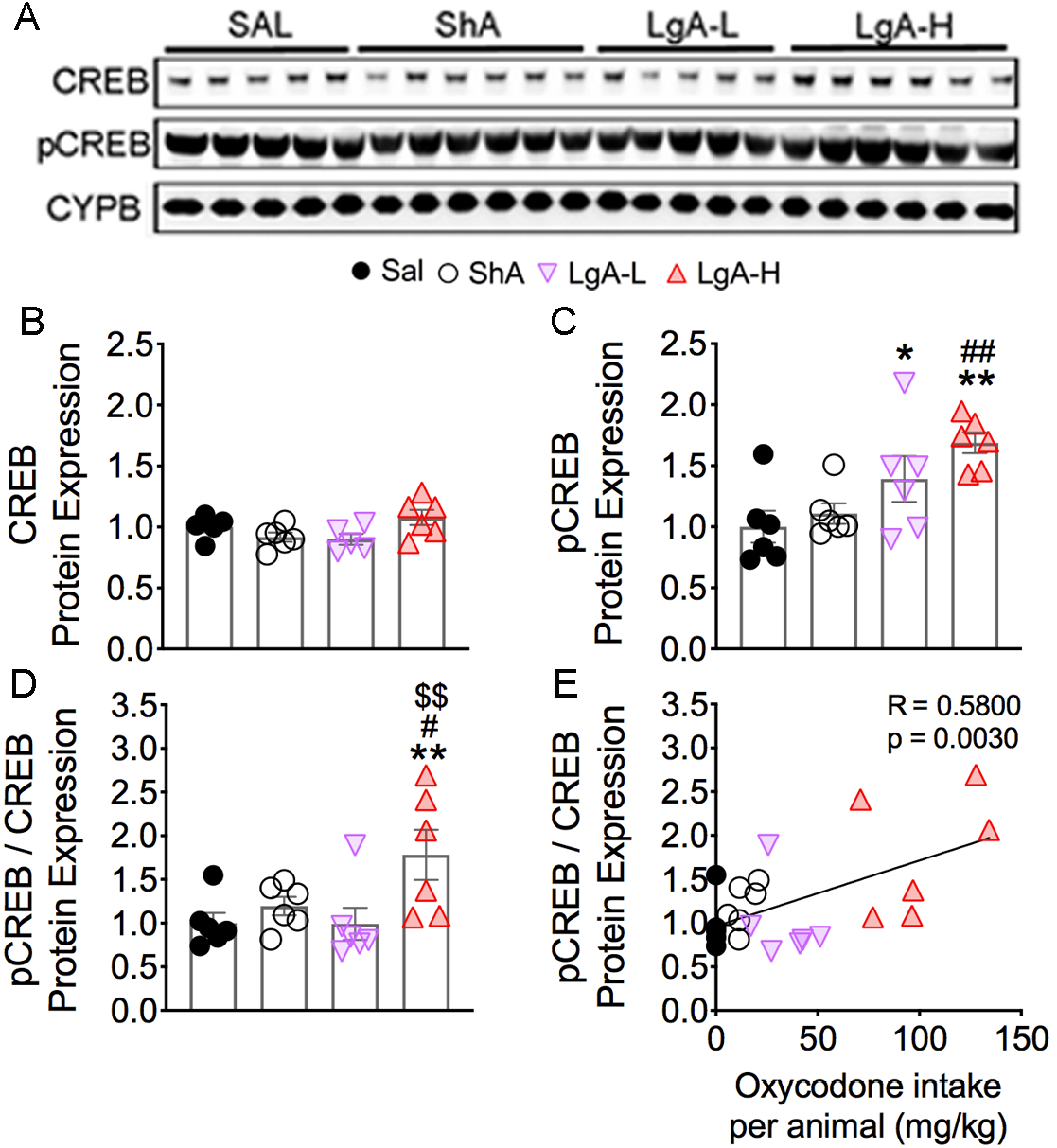
Increased phosphorylation of CREB protein levels in the LgA-H rats. **(A)** Images of western blot and **(B, C)** quantification of CREB and pCREB. **(B)** CREB protein levels show no significant changes. **(C)** pCREB is significantly increases in LgA-L and LgA-H rats. **(D)** pCREB/CREB ratios are increased in the LgA-H rats and **(E)** correlated with amount of oxycodone (n=5-6 Sal; n=6 ShA; n=5-6 LgA-L; n=6 LgA-H). Key to statistics: *, ** = p < 0.05, 0.01, respectively, in comparison to Sal rats; #, ## = p < 0.05, 0.01, respectively, in comparison to SHA rats; $$ = p < 0.01 in comparison to LgA-L rats. Statistical analyses are as described in Fig. 2.

### Oxycodone SA induces increased phosphoacetylation in LgA rats

In addition to CREB phosphorylation, activated MSK1 and MSK2 participate in the phosphorylation of H3 at serine residue S10 and can cause increases in H3 phosphoacetylation of H3S10pK14Ac ^34,37,38^. We therefore sought to determine the effects of oxycodone SA on the abundance of this histone marker. Figure 5 shows the results for H3 and H3S10pK14Ac. Unexpectedly, we found significant decreases [F_(3, 20)_ = 5.02, p = 0.0094] in H3 protein levels in the ShA group in comparison with Sal and the LgA-H groups (Figs. 5A, 5B). In contrast, H3S10pK14Ac abundance was significantly increased [F_(3, 20)_ = 20.0, p < 0.0001] in the LgA-L and LgA-H groups compared with Saline rats. Moreover, H3S10pK14Ac in the LgA-H group was substantially increased compared to the other 3 groups (Figs. 5A, 5C). H3S10pK14Ac/H3 ratios were also increased [F_(3, 20)_ = 20.0, p < 0.0001] in the LgA-L and LgA-H groups compared to the Saline and ShA groups (Fig. 5D). Regression analyses revealed that both H3S10pK14Ac abundance and H3S10pK14Ac/H3 ratios positively correlated with the amount of total oxycodone taken during the experiment (Figs. 5E, 5F).

**Figure 5.**
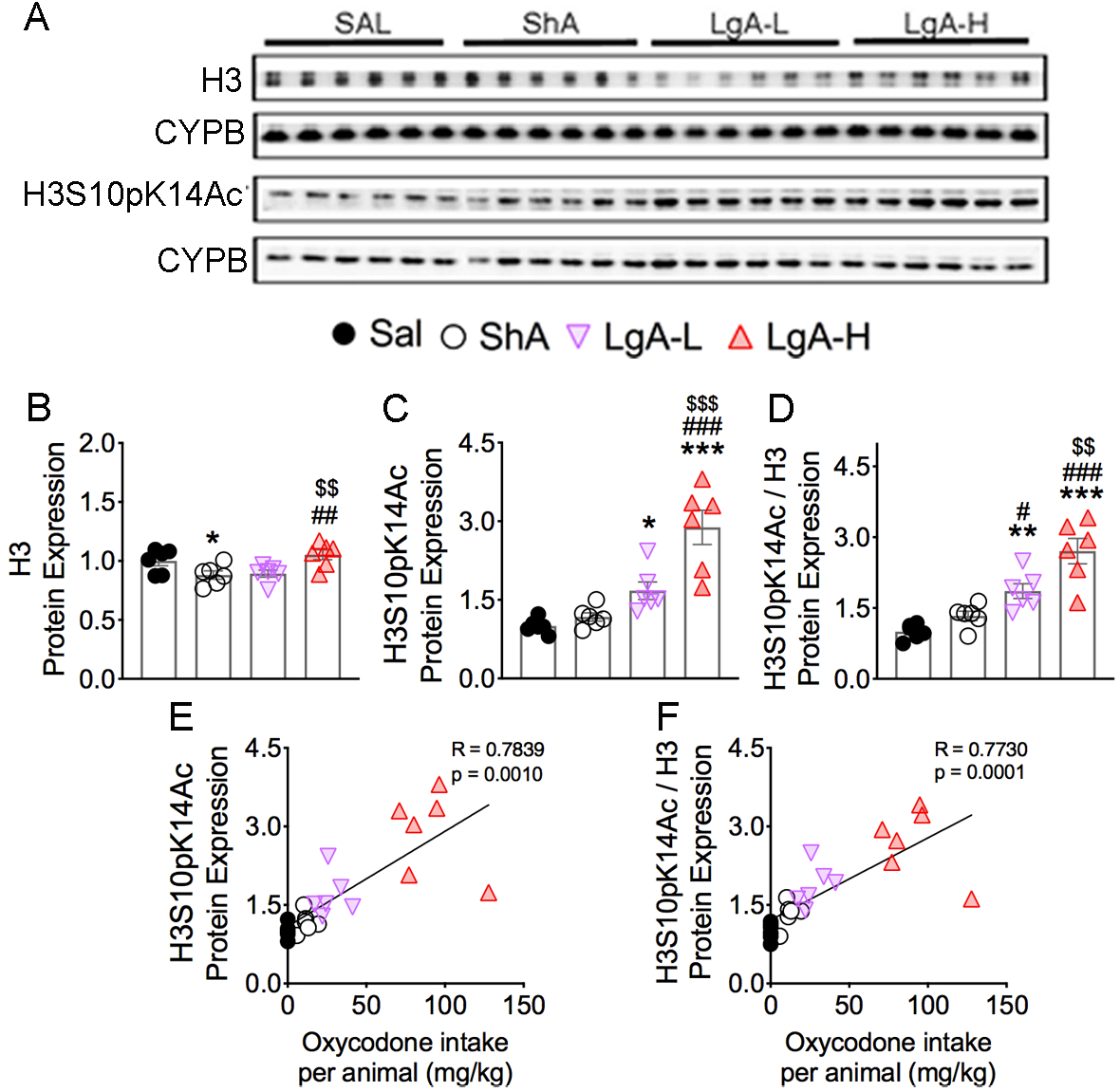
Effects of oxycodone SA and early withdrawal on H3 and H3S10pK14Ac. **(A)** Images of western blot and quantification of **(B)** H3 and **(C)** H3S10pK14Ac protein levels. **(B)** Protein levels of H3 display decrease in the ShA group. **(C)** Protein expression of H3S10pK14Ac shows significant increases in LgA-L and LgA-H groups. **(D)** Ratio of H3S10pK14Ac/H3 displays significant increases in the LgA-L and LgA-H groups. **(E)** Protein expression of H3S10pK14Ac and **(F)** H3S10pK14Ac/H3 positively correlated with doses of oxycodone taken during the experiment (n=5-6 Sal; n=6 ShA; n=6 LgA-L; n=6 LgA-H). Key to statistics: *, **, *** = p < 0.05, 0.01, 0.001, respectively, in comparison to Sal rats; #, ##, ### = p < 0.05, 0.01, 0.001, respectively, in comparison to SHA rats; $$, $$$ = p < 0.01, 0.001 in comparison to LgA-L rats. Statistical analyses are as described in Fig. 2.

### Differential protein expression of CBP and H3K27Ac in oxycodone exposed rats

Phosphorylated CREB recruits CBP, under certain circumstances, to promote changes in gene expression ^39,40^. We thus tested the possibility that oxycodone SA might have influenced striatal CBP protein expression and found that CBP protein expression was significantly increased [F_(3, 17)_ = 6.82, p = 0.0032] in the LgA-L and LgA-H rats compared with the Saline and ShA groups (Figs. 6A, 6B). Increased CBP expression correlated with the amount of oxycodone consumed by the rats (Fig. 6C).

**Figure 6.**
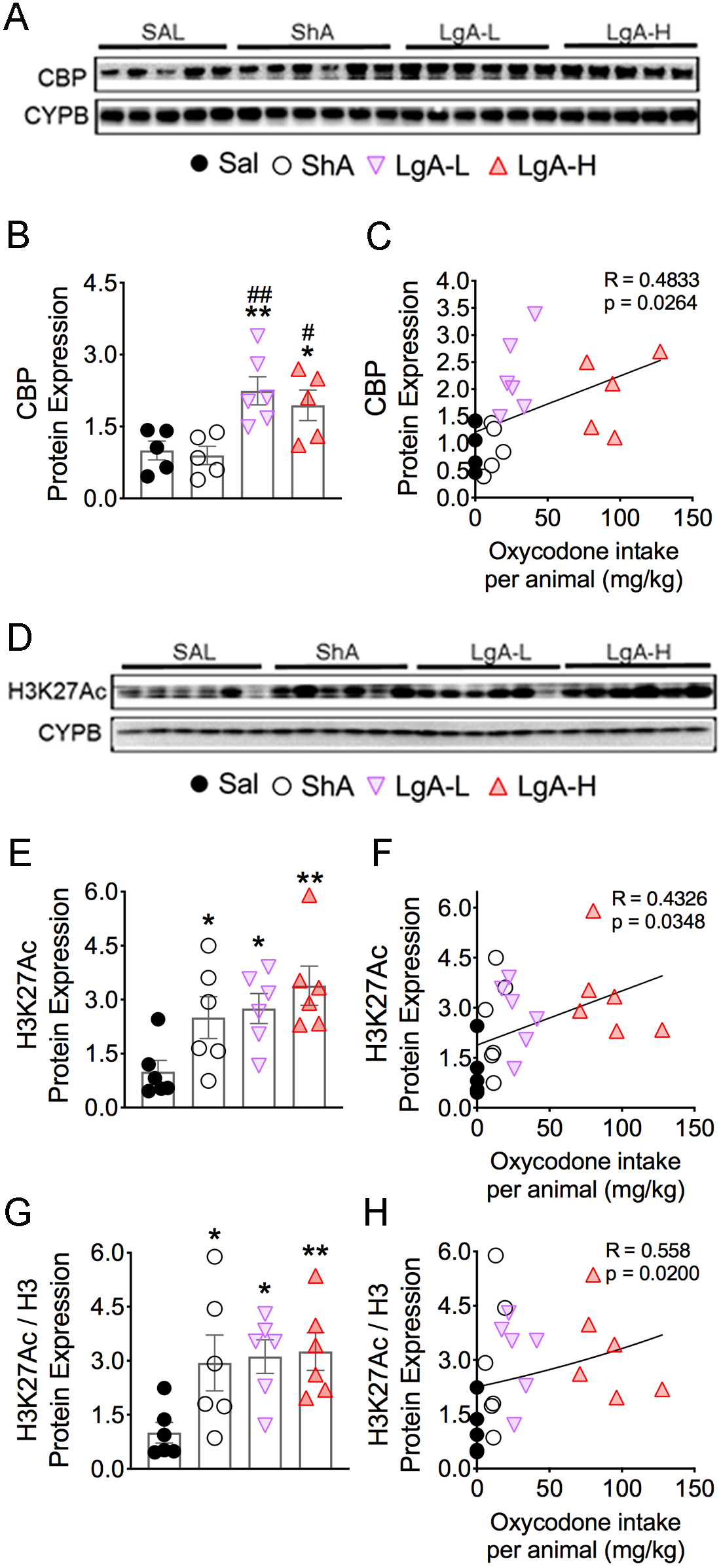
Differential effects on the protein expression of CBP and H3K27Ac after oxycodone SA. **(A, D)** Images of western blot and quantification of **(B)** CBP and **(E)** H3K27Ac protein levels. **(B)** CBP protein levels display significant increases in LgA groups. The protein levels of **(E)** H3K27Ac and **(G)** H3K27Ac/H3 protein were significantly increased in all drug groups. The regression analyses of **(C)** CBP, **(F)** H3K27Ac, and **(H)** H3K27Ac/H3 correlates with the amount of oxycodone taken (n=5-6 Sal; n=5-6 ShA; n=6 LgA-L; n=5-6 LgA-H). Key to statistics: *, ** = p < 0.05, 0.01, respectively, in comparison to Sal rats; #, ## = p < 0.05, 0.01, respectively, in comparison to ShA rats. Statistical analyses are as described in Fig. 2.

In addition to CBP’s transcriptional co-activity, it functions as a histone acetyltransferase ^41,42^ that mediates acetylation of H3K27 ^43–45^, a marker of active enhancers ^46–48^ that is involved in regulating neuronal gene expression. We therefore measured the abundance of H3K27Ac, which had been previously shown to be impacted in the brains of heroin addicts ^49^. We found significant increases [F_(3, 20)_ = 4.53, p = 0.0140] in H3K27Ac abundance in all oxycodone groups, including ShA rats that did not escalate their intake (Figure 6D, 6E), suggesting that oxycodone exposure is enough to increase striatal H3K27Ac abundance. We found that H3K27Ac abundance positively correlated with the amount of oxycodone self-administered by rats (Fig. 6F). These increases confirm the data in heroin-using individuals ^49^. H3K27Ac/H3 ratios were also significantly increased [F_(3, 20)_ = 3.82, p = 0.0259] in the 3 oxycodone groups (Fig. 6G) and correlated with the amount of oxycodone taken (Fig. 6H).

Because CBP expression was only increased in the two LgA groups while H3K27Ac was increased in the 3 oxycodone groups, we sought to determine if the expression of another histone H3 acetyltransferase, Tip60, with putative activity towards the lysine 27 residue ^50^ was affected in the 3 oxycodone groups. Indeed, Tip60 protein expression was significantly increased [F_(3, 20)_ = 3.63, p = 0.0307] in all three oxycodone groups (Supplementary Fig. S4).

### Oxycodone SA increases *GluA2* and *GluA3* glutamate receptor mRNA levels

Changes in gene expression in response to exogenous stimuli include many target genes that are expressed with different time courses of induction. Because respective changes in histone phosphorylation and acetylation generated by MSKs and CBP are known regulators of gene expression ^51–54^, we tested the idea that some of their target genes might be affected in the striata of oxycodone-exposed rats. We also measured the mRNA levels of some glutamatergic genes whose expression was altered in the brains of heroin users based on a previous microarray study ^49^. Figure 7 shows the effects of oxycodone SA on the mRNA expression of *GluA1*, *GluA2*, *GluA3*, and *GluA4* subunits of AMPA receptors ^55^. We found no substantial changes in *GluA1* [F_(3, 29)_ = 2.03, p = 0.1322; Fig. 7A] and *GluA4* [F_(3, 34)_ = 1.60, p = 0.2084; Fig. 7G] mRNA levels, with no relationships between their levels and the amount of oxycodone consumed during the experiment (Figs. 7B, 7H). In contrast, striatal *GluA2* [F_(3, 27)_ = 3.49, p = 0.0291; Fig. 7C] and *GluA3* [F_(3, 31)_ = 15.7, p < 0.0001; Fig. 7E] were increased in the LgA-H group compared to other groups. Regression analyses revealed significant oxycodone amount-dependent changes in their mRNA levels (Figs. 7D, 7F). Moreover, the changes in *GluA2* and *GluA3* mRNAs correlated with changes in pMSK1 (Figs. 8A, 8B). However, only changes in *GluA3*, but not in *GluA2*, mRNA levels correlated with changes in pMSK2, H3S10pK14Ac, and CBP protein expression (Figs. 8C-8H).

**Figure 7.**
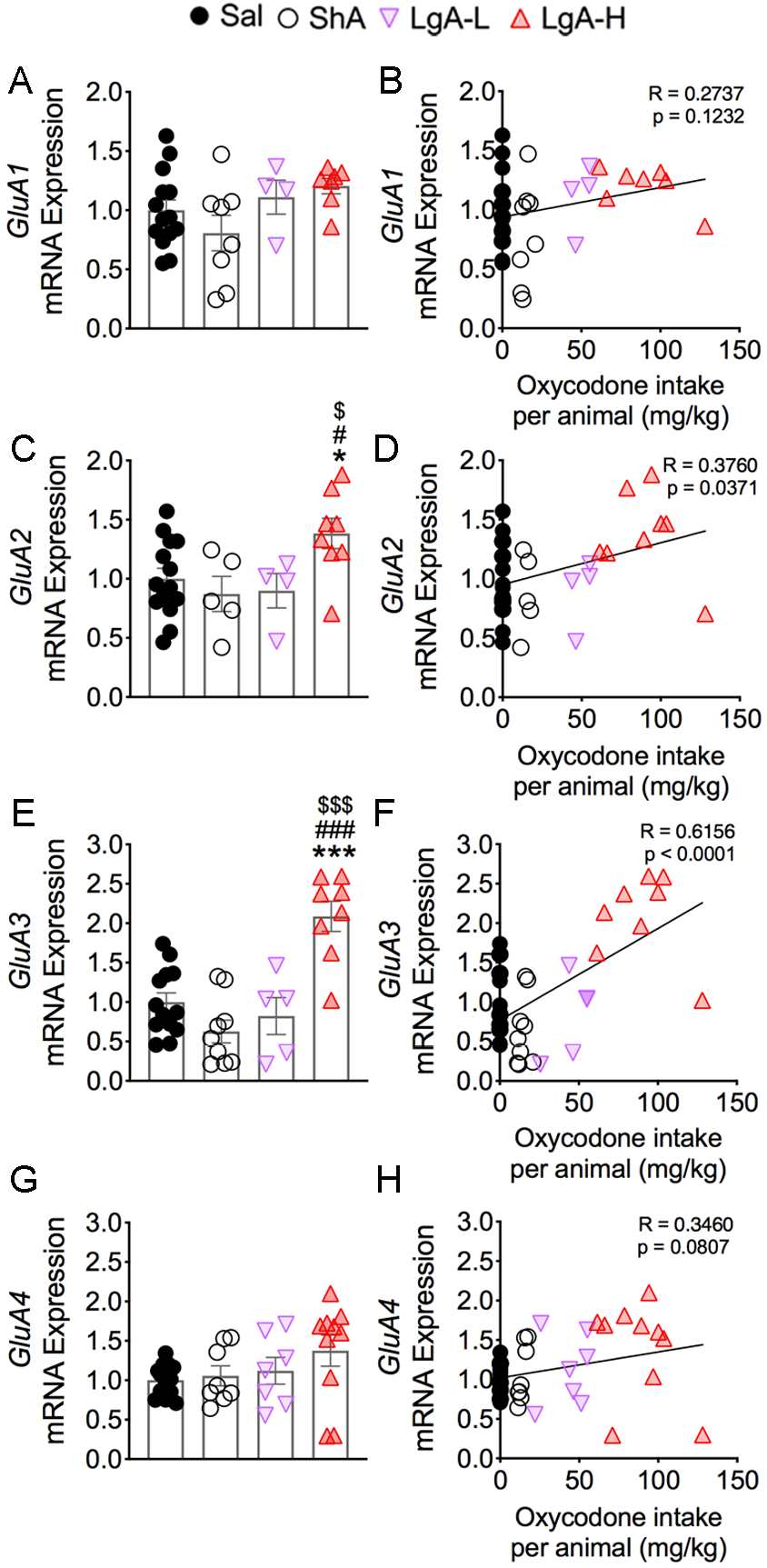
Consequences of oxycodone SA and early withdrawal on mRNA on *GluA1*, *GluA2*, *GluA3*, and *GluA4* mRNA levels. mRNA levels of **(A)** *GluA1* and **(G)** *GluA4* were not significantly affected. There were no correlations between **(B)** *GluA1* and **(H)** *GluA4* mRNA levels and the consumption of oxycodone. However, (**C**) *GluA2* and (**E**) *GluA3* mRNA levels were significantly increased in the LgA-H rats, with significant positive correlations between mRNA expression of **(D)** *GluA2* and **(F)** *GluA3* with the amount of oxycodone taken. Key to statistics: *, *** = p < 0.05, 0.001, respectively, in comparison to Sal rats; #, ### = p < 0.05, 0.001, respectively, in comparison to SHA rats; $, $$$ = p < 0.05, 0.001 in comparison to LgA-L rats (n=10-11 Sal; n=5-9 ShA; n=4-7 LgA-L; n=7-10 LgA-H). Statistical analyses are as described in Fig. 2.

**Figure 8.**
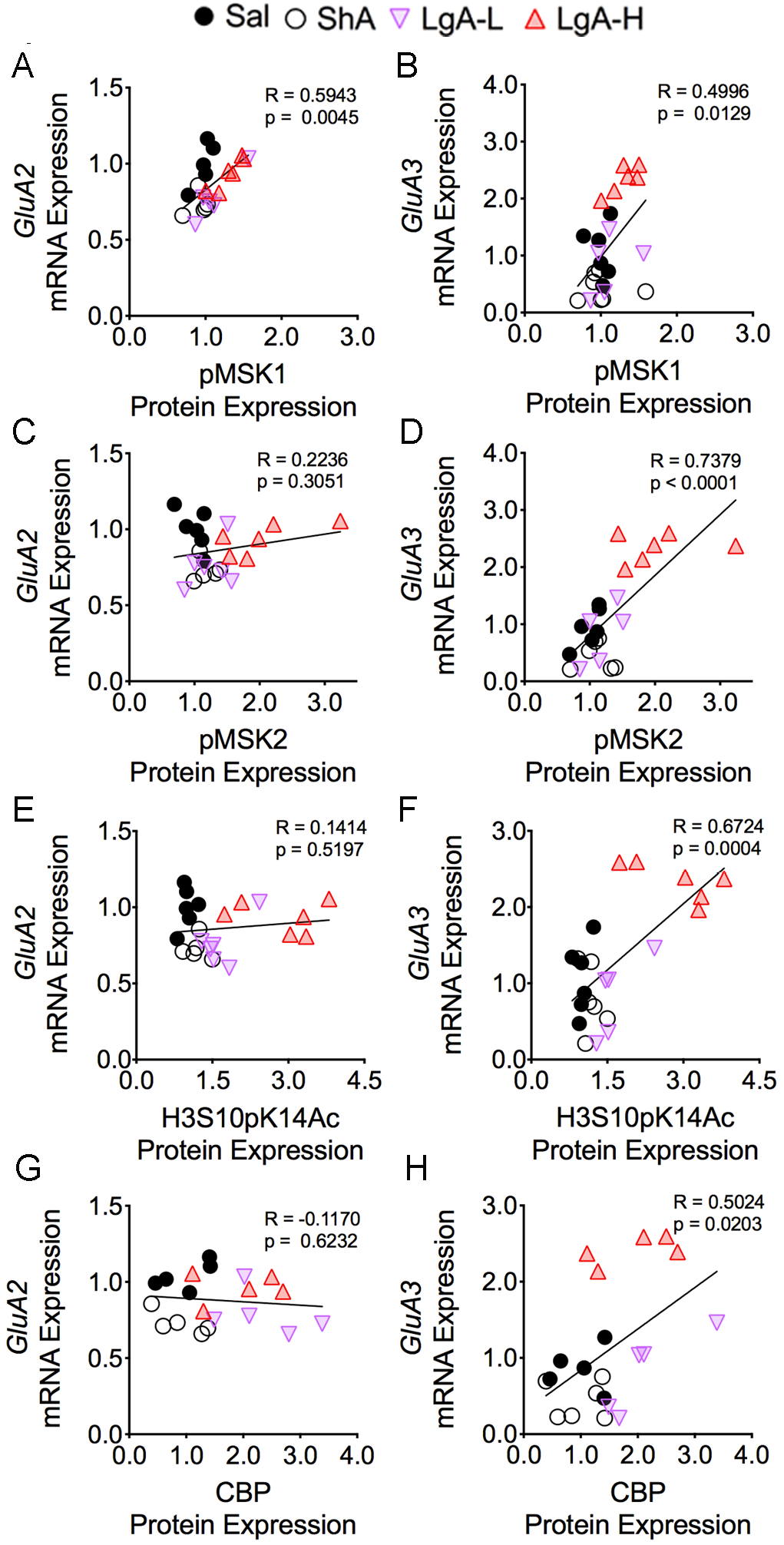
*GluA3* mRNA correlated with pMSK1, pMSK2, H3S10pK14Ac, and CBP protein expression. The mRNA expression of *GluA2* shows significant linear relationship with **(A)** pMSK1, but not with **(C)** pMSK2, **(E)** H3S10pK14Ac, and **(G)** CBP protein expression. However, the mRNA levels of *GluA3* display a positive correlation with the protein abundance of **(B)** pMSK1, **(D)** pMSK2, **(F)** H3S10pK14Ac, and **(H)** CBP (n=5-6 Sal; n=5-6 ShA; n=5-6 LgA-L; n=5-6 LgA-H). The correlation coefficients and p values are shown on the graph.

### mRNA levels of acetyltransferases, *Ncoa1*-3, are increased in LgA oxycodone rats

Egervari et al. (2017) had recently reported that the expression of the acetyltransferase, *nuclear receptor coactivator 1* (*Ncoa1*), was significantly increased in the ventral striatum of heroin users. We therefore measured the expression of *Ncoa1*-3 in our experiment. Supplemental Figure 3 shows significant increases in the mRNA levels of *Ncoa1* [F_(3, 35)_ = 12.6; p < 0.0001], *Ncoa2* [F_(3, 31)_ = 4.55; p = 0.0094], and *Ncoa3* [F_(3, 28)_ = 12.5; p < 0.0001] in the LgA-H group compared to the other groups. Furthermore, there were significant correlations between their expression and changes in pMSK1, pMSK2, and H3S10pK14Ac protein abundance (Supplementary Fig. S3). However, we observed no significant correlations between *Ncoa1*, *Ncoa2*, and *Ncoa3* mRNA levels and CBP suggesting that histone phosphorylation might play a more important role in regulating their expression than histone acetylation (Supplementary Fig. S3).

## Discussion

In the present study, we show that long-access to oxycodone SA over a period of 20 days leads to activation of several kinases involved in the MAPK/MSK signaling pathway with consequent CREB and histone H3 phosphorylation in the rat dorsal striatum. These results are consistent, in part, with previous evidence of the involvement of the MAPK in the biochemical effects of morphine ^56,57^. We also found significant increases in CBP and H3K27 acetylation in oxycodone-exposed rats. These findings are consistent with observations that epigenetic mechanisms are involved in models of opioid abuse ^58^. The changes in signaling pathways are accompanied by oxycodone-induced increased gene expression of *GluA2* and *GluA3* subunits of AMPA receptors and of the acetyltransferases, *Ncoa1*-3. Our results provide novel insights into the role of H3S10pK14Ac in oxycodone-induced gene expression and hint to a model whereby this histone marker is involved in the regulation of genes that might be responsible for some long-term molecular adaptations that drive compulsive oxycodone intake.

The dorsal striatum is a brain region that is integral to various behavioral changes consequent to drug taking behaviors including habit forming and drug seeking during periods of opioid withdrawal ^19,22,23,59^. Similar to other observations with cocaine, methamphetamine, and other drugs ^58,60–66^, we found increased phosphorylation of PKC, ERK1/2, MSK1/2, and pCREB in the rat dorsal striatum after repeated exposure to oxycodone SA. Increased phosphorylation of H3S10pK14Ac, a marker that is downstream of these kinases ^34,37,67^ is of interest because these findings suggest that repeated exposure to long-access oxycodone self-administration might engender a permissive molecular state characterized by increased histone H3 phosphoacetylation and a more open chromatin structure. The hypothesized permissive state might also facilitate pCREB binding at the cAMP-Response Element (CRE) on the promoters of genes that have been implicated in the regulation of synaptic plasticity ^51,68,69^. CREB activation is also known to enhance the recruitment of co-activators ^39,40^ such as CBP, an acetyltransferase that acetylates H3K27 ^43–45,70^ to increase the transcription of downstream genes in diverse cell populations ^39,71,72^. This suggestion is further supported by observations of increased expression of co-activators for the steroid hormone receptor family, *Ncoa1* ^73^, *Ncoa2* ^74^, and *Ncoa3* ^75^, that can enhance transcription, in part, via histone acetylation ^76,77^ and recruitment of CBP ^75,78,79^, whose expression is also increased in rats exposed to relatively large quantities of oxycodone. Our proposal of an oxycodone-induced permissive state in the dorsal striatum is consistent with observations that histone H3 phosphorylation and acetylation can work in concert to regulate gene expression ^80^. This discussion is supported by the observations that H3 phosphoacetylation also participates in heroin-induced conditioned place preference ^81^, thus indicating a role of phosphoacetylation in the effects of opioids in general.

Our observations of increased H3 phosphorylation and acetylation led us to test the possibility that some genes downstream of these molecular events might show differential expression in the brains of oxycodone-exposed rats. Indeed, we found increased expression of several genes in the LgA-H rats that showed increased abundance of striatal pMSK1, pMSK2, and histone H3S10pK14Ac. Of interest among those are the changes in AMPA receptor subunits, *GluA2* and *GluA3,* in LgA-H rats in an oxycodone amount- and pMSK1-dependent fashion. MSK1 and MSK2 are known to play substantial roles in a number of biological events including synaptic plasticity ^82^. The altered expression of *GluA2* and *GluA3* is of singular interest because Egervari et al. (2017) had reported that their microarray analyses, using tissues from the ventral striatum of heroin users, had detected changes in the expression of several genes, including *GluA3*, which are involved in glutamate neurotransmission. Increases in the expression of *GluA2* and *GluA3* receptor subunits in our study are consistent with the ^83,84^proposed roles of glutamate receptors in substance use disorder includes cocaine ^85^, methamphetamine ^86^, and opioids ^83,84^. For example, chronic cocaine increases *GluA2* expression in the nucleus accumbens and increased expression of *GluA2* via viral injections enhanced the sensitivity of mice to the behavioral effects of cocaine ^85^, thus suggesting that increased *GluA2* expression in the present study might have served to facilitate escalation of oxycodone intake in the LgA-H rats. A similar argument could be made for our novel findings of increased *GluA3* expression after oxycodone SA. *GluA3*- containing AMPA receptors are located in various brain regions ^87,88^. Because *GluA3* exists in *GluA2*A3 combinations or *GluA3* monomers or dimers ^89^, it is possible that increased expression of both *GluA2* and *GluA3* might potentiate AMPAR-mediated changes in synaptic plasticity during repeated oxycodone exposure. Alternatively, *GluA3* alone may regulate oxycodone intake because *GluA3* knockout mice show decreased alcohol intake ^90^. Thus, elucidation of the specific roles that *GluA3* alone or in combination with *GluA2* play in oxycodone SA will await future genetic and pharmacological studies.

In conclusion, we have demonstrated that rats that self-administer large quantities of oxycodone showed increased histone and CREB phosphorylation via activation of the MAPK/MSK phosphorylation signaling pathway in the rat dorsal striatum. Rats exposed to large quantities of oxycodone also showed increased striatal CBP and histone acetylation in oxycodone-exposed rats. Changes in histone modifications are proposed to lead to more permissive chromatin states that promoted changes in the expression in a diversity of classes of genes as exemplified by increased mRNA levels of acetyltransferases, *Ncoa1*-3, and of *AMPA* receptor subunits, *GluA2* and *GluA3*, in an oxycodone amount-dependent fashion. Importantly, changes in the expression of both *GluA2 and GluA3 mRNA* levels correlated with altered abundance of pMSK1, a kinase that is involved in the regulation of synaptic plasticity ^91^. These suggestions are illustrated schematically in Figure 9. Finally, involvement of MAPK/MSK/histone phosphorylation in oxycodone SA identifies these kinases as potential pharmacological targets against oxycodone use disorder.

**Figure 9.**
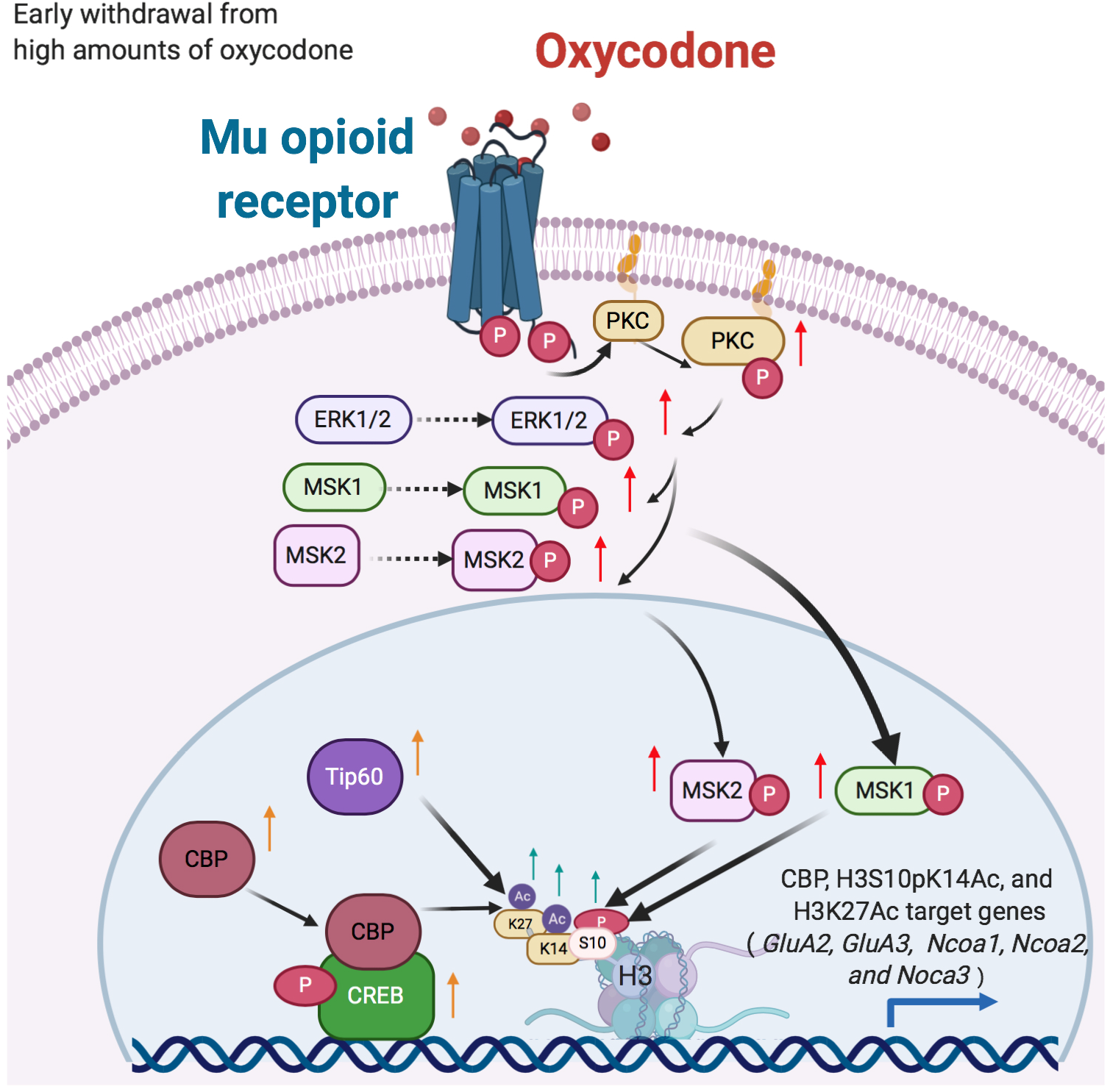
Illustration of the activation of MAPK/MSK signaling pathway in LgA-H rats. Intake of large amount of oxycodone during long-access over 20 days caused increased phosphorylation of PKC, ERK1/2, MSK1, and MSK2. Increased MSK phosphorylation is accompanied by increased histone H3 phosphoacetylation and CREB phosphorylation. In addition, there were increases in the protein expression of two acetyltransferases, CBP and Tip60 that acetylate H3K27. Recruitment of CBP by CREB and histone modifications serve to create a permissible transcriptional environment that led to increased mRNA levels of *GluA2* and *GluA3* in a MSK1-dependent fashion. This kinase-histone modification cascade may serve as targets for therapeutic interventions against oxycodone use disorder.

## Author contributions

C.A.B, M.T.M, and B.L performed self-administration, western blot and RT-qPCR experiments. C.A.B and J.L.C prepared manuscript. J.L.C supervised the overall project.

---

**Supplementary Figure 1.**
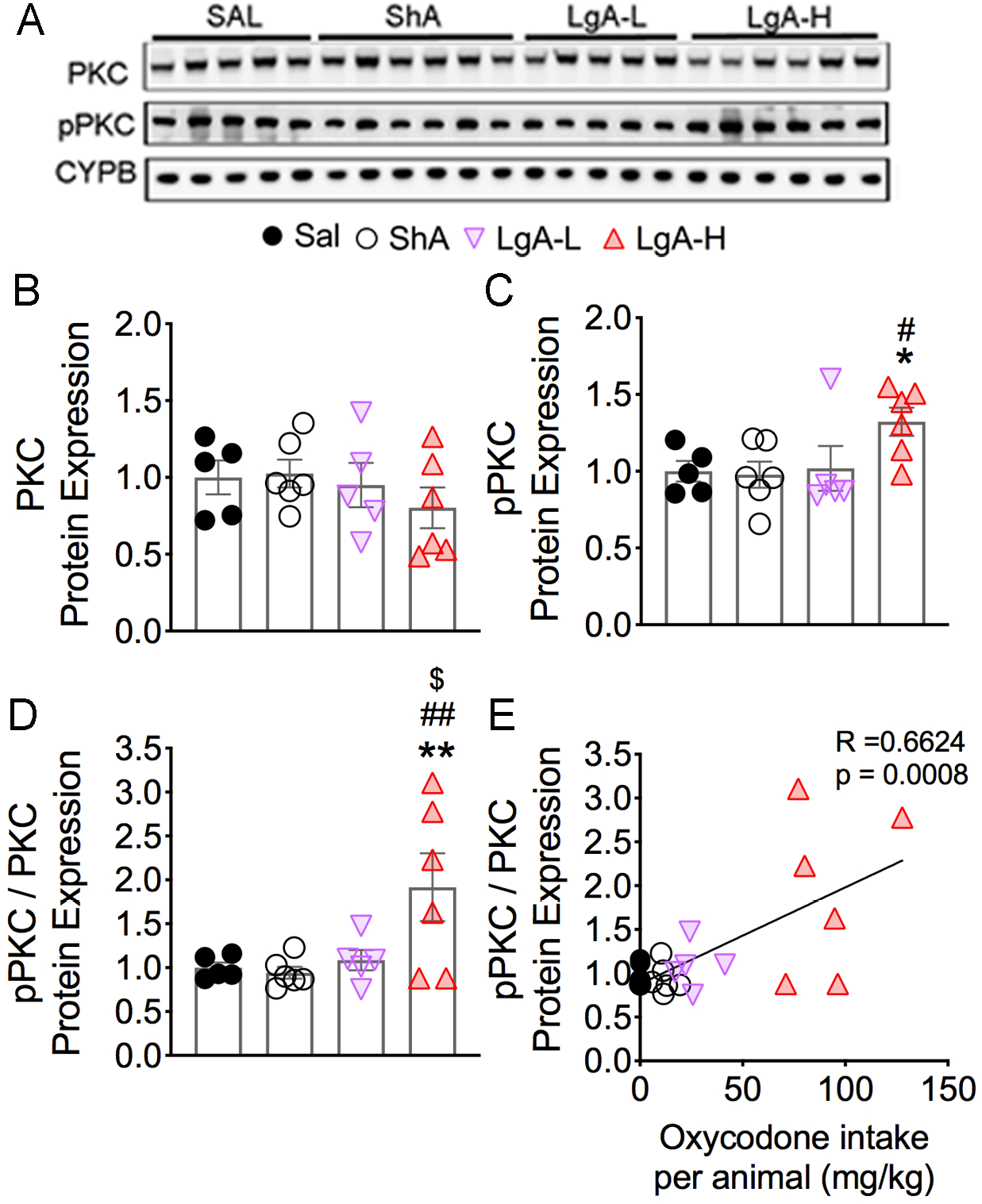
Effects of oxycodone SA on PKC phosphorylation. **(A)** Images of western blot and **(B, C)** quantification of PKC and pPKC. **(B)** PKC protein levels were not significantly affected in any group. **(C)** pPKC abundance is increased in only LgA-H rats. **(D)** pPKC/PKC ratios are increased in only the LgA-H rats. **(E)** pPKC/PKC ratios correlate with amount of oxycodone taken during the experiment (n=5-6 Sal; n=6 ShA; n=5 LgA-L; n=6 LgA-H). Key to statistics: *, ** = p < 0.05, 0.01, respectively, in comparison to Sal rats; #, ## = p < 0.05, 0.01, respectively, in comparison to ShA rats; $ = p < 0.05 in comparison to LgA-L rats. Statistical analyses were performed by one-way ANOVA followed by Bonferroni or Fisher’s PLSD post hoc test. The correlation coefficients and p values are shown on the graph.

**Supplementary Figure 2.**
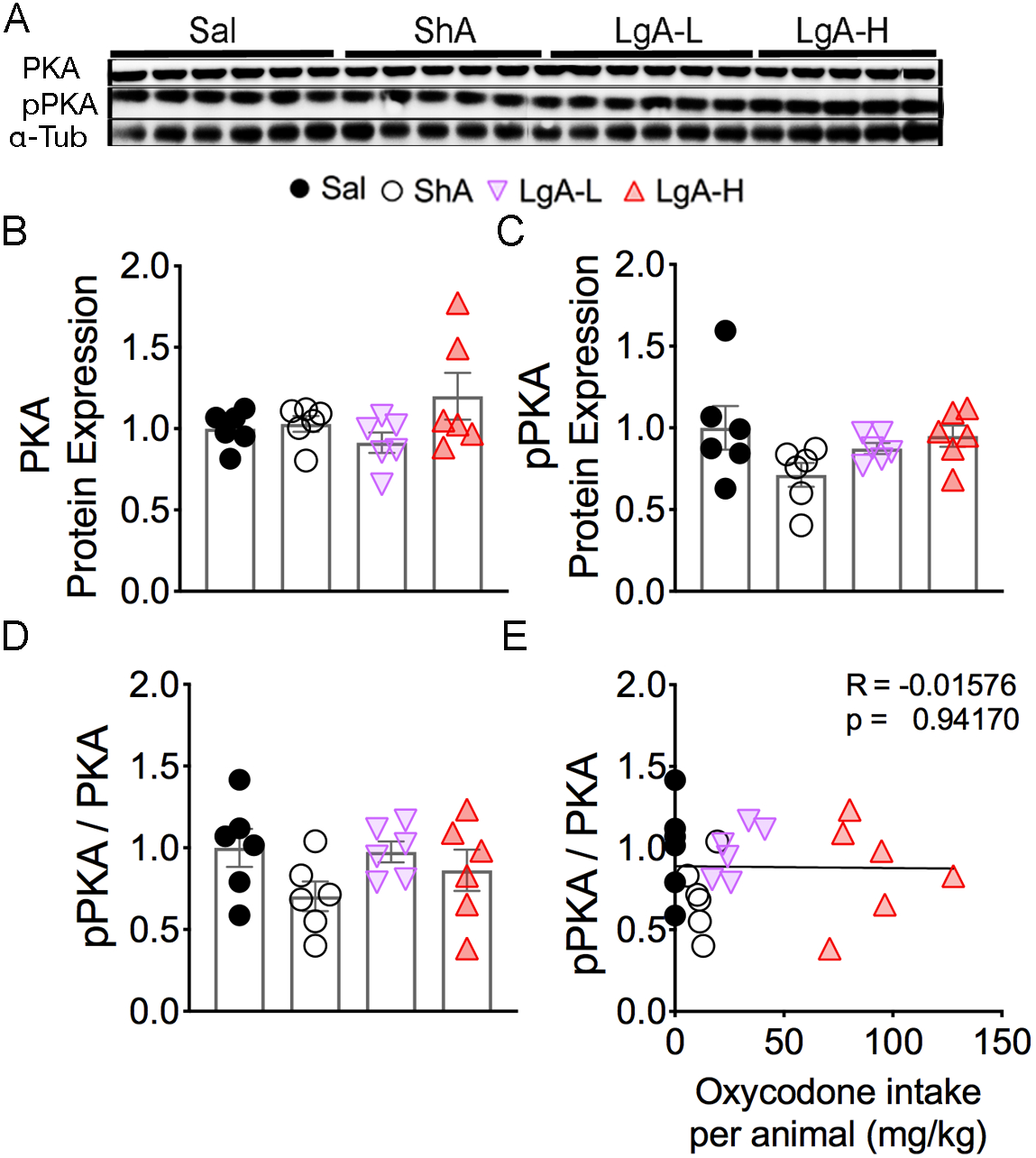
Exposure to oxycodone SA shows no changes in PKA. **(A)** Images of western blot and quantification of **(B)** PKA and **(C)** pPKA protein expression. Protein levels of **(B-D)** PKA, pPKA, and pPKA / PKA ratios show no significant changes. Similarly, **(E)** ratio of pPKA / PKA showed no correlation to the amount of oxycodone taken (n=6 Sal; n=6 ShA; n=6 LgA-L; n=6 LgA-H). Statistical analyses were performed by one-way ANOVA followed by Bonferroni or Fisher’s PLSD post hoc test. The correlation coefficients and p values are shown on the graph.

**Supplementary Figure 3.**
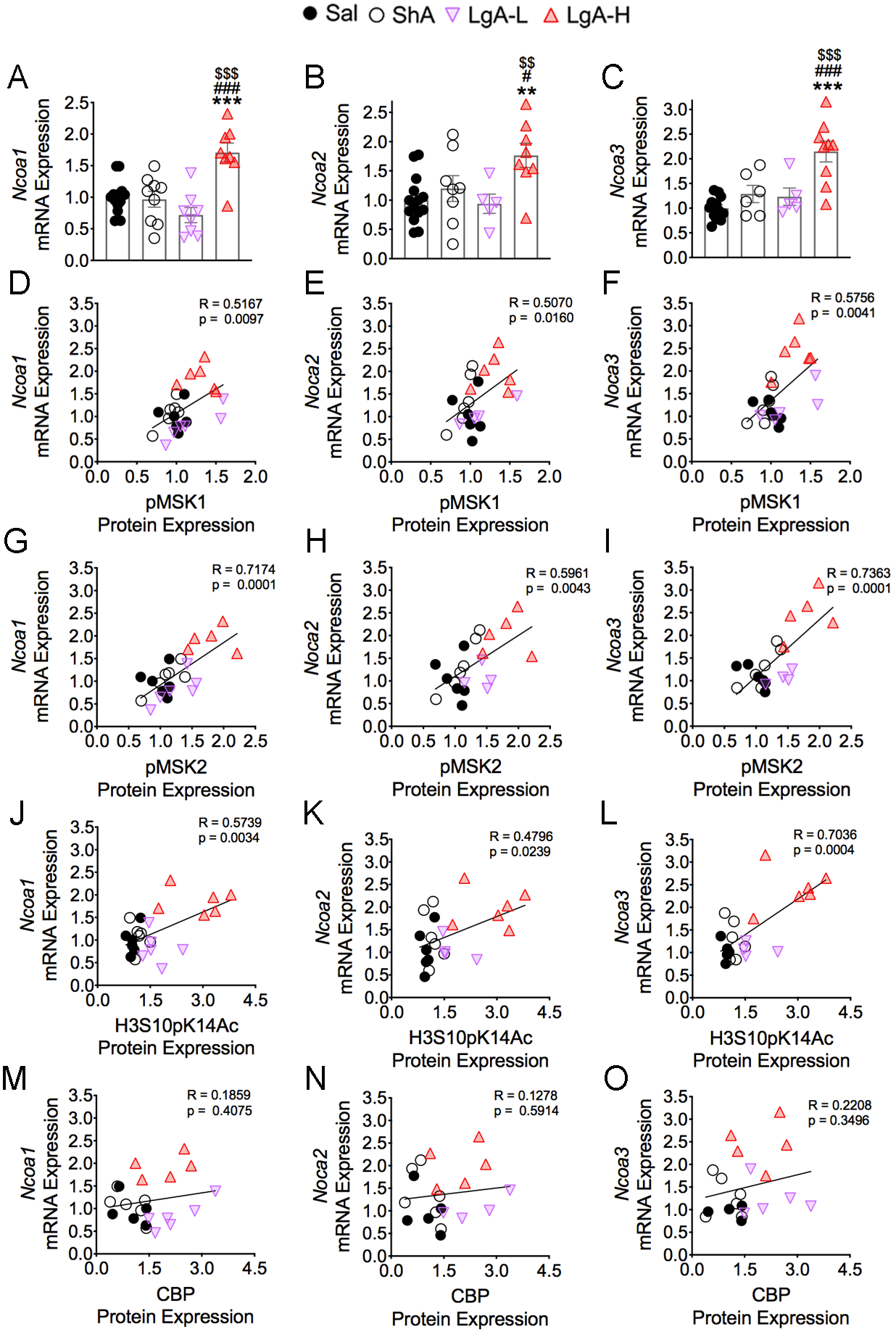
mRNA expression on the members of Ncoa family after exposure to oxycodone SA and early withdrawal. The mRNA levels of **(A)** *Ncoa1*, **(B)** *Ncoa2*, and **(C)** *Ncoa3* show significant increases in the LgA-H group. The mRNA levels of *Ncoa1*, *Ncoa2*, and *Ncoa3* show a positive correlation with **(D-F)** pMSK1, **(G-I)** pMSK2, and **(J-L)** H3S10pK14Ac. However, the mRNA expression of *Ncoa1, Ncoa2, and Ncoa3* shows no correlation **(M-O)** CBP protein expression (n=4-9 Sal; n=5-9 ShA; n=4-8 LgA-L; n=5-9 LgA-H). Key to statistics: **, *** = p < 0.01, 0.001, respectively, in comparison to Sal rats; #, ### = p < 0.05, 0.001, respectively, in comparison to SHA rats; $$, $$$ = p < 0.01, 0.001, respectively, in comparison to LgA-L rats. Statistical analyses are as described in Fig. 2.

**Supplementary Figure 4.**
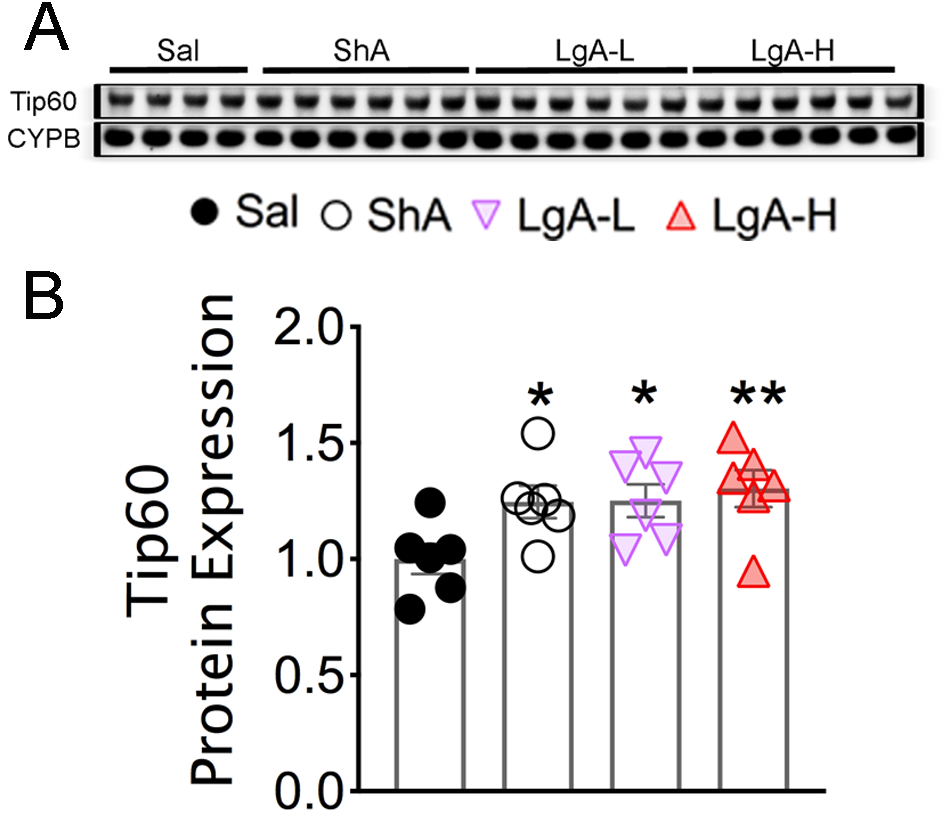
Exposure to oxycodone SA increases Tip60. **(A)** Images of western blot and **(B)** quantification of Tip60 protein levels show increases in all drug groups (n=5 Sal; n=6 ShA; n=6 LgA-L; n=6 LgA-H). Key to statistics: *, **= p < 0.05, 0.01, respectively, in comparison to Sal rats. Stats were performed by one-way ANOVA followed by Bonferroni or Fisher’s PLSD post hoc test.

**Supplementary Table 1.** Antibody list

| Symbol | Source | Dilution | Catalog # | RRID | Company |
| --- | --- | --- | --- | --- | --- |
| PKC | Rabbit | 1:1,000 | 2056 | AB2284227 | Cell Signaling, Danvers, MA, USA |
| pPKC | Mouse | 1:1,000 | SC-377565 | AB10611031 | Santa Cruz Bio. Tech., Dallas, TX, USA |
| ERK1/2 | Rabbit | 1:1,000 | 06-182 | AB310068 | Sigma-Aldrich, St. Louis, MO, USA |
| pERK1/2 | Rabbit | 1:1,000 | 4370 | AB2315112 | Cell Signaling, Danvers, MA, USA |
| MSK1 | Rabbit | 1:1,000 | 3489 | AB2285349 | Cell Signaling, Danvers, MA, USA |
| pMSK1 | Rabbit | 1:1,000 | 9595 | AB2181783 | Cell Signaling, Danvers, MA, USA |
| MSK2 | Rabbit | 1:10,000 | 3679 | AB2181641 | Cell Signaling, Danvers, MA, USA |
| pMSK2 | Rabbit | 1:10,000 | SAB4504629 | AB11184854 | Sigma-Aldrich, St. Louis, MO, USA |
| CREB | Rabbit | 1:1,000 | 9197 | AB331277 | Cell Signaling, Danvers, MA, USA |
| pCREB | Mouse | 1:1,000 | 9196 | AB331275 | Cell Signaling, Danvers, MA, USA |
| H3 | Rabbit | 1:10,000 | 06-755 | AB2118461 | Sigma-Aldrich, St. Louis, MO, USA |
| H3S10pK14Ac | Rabbit | 1:1,000 | 07-081 | AB310366 | Sigma-Aldrich, St. Louis, MO, USA |
| H3K27Ac | Rabbit | 1:1,000 | 39034 | AB2722569 | Active Motif, Carlsbad, CA, USA |
| CBP | Rabbit | 1:500 | 7389 | aB2616020 | Cell Signaling, Danvers, MA, USA |
| PKA | Mouse | 1:1,000 | 4782 | AB2170170 | Cell Signaling, Danvers, MA, USA |
| pPKA | Rabbit | 1:1,000 | 5661 | AB10707163 | Cell Signaling, Danvers, MA, USA |
| TIP60 | Rabbit | 1:10,000 | MABE430 | AB2551948 | Sigma-Aldrich, St. Louis, MO, USA |
| CYPB | Rabbit | 1:10,000 | ab16045 | AB443295 | Abcam, Cambridge, MA, USA |
| Alpha-Tubulin | Mouse | 1:25,000 | T6074 | AB477582 | Sigma-Aldrich, St. Louis, MO, USA |
| HRP (rabbit) | Goat | 1:5,000 | 7074 | AB2099233 | Cell Signaling, Danvers, MA, USA |
| HRP (mouse) | Horse | 1:5,000 | 7076 | AB330924 | Cell Signaling, Danvers, MA, USA |

**Supplementary Table 2.**
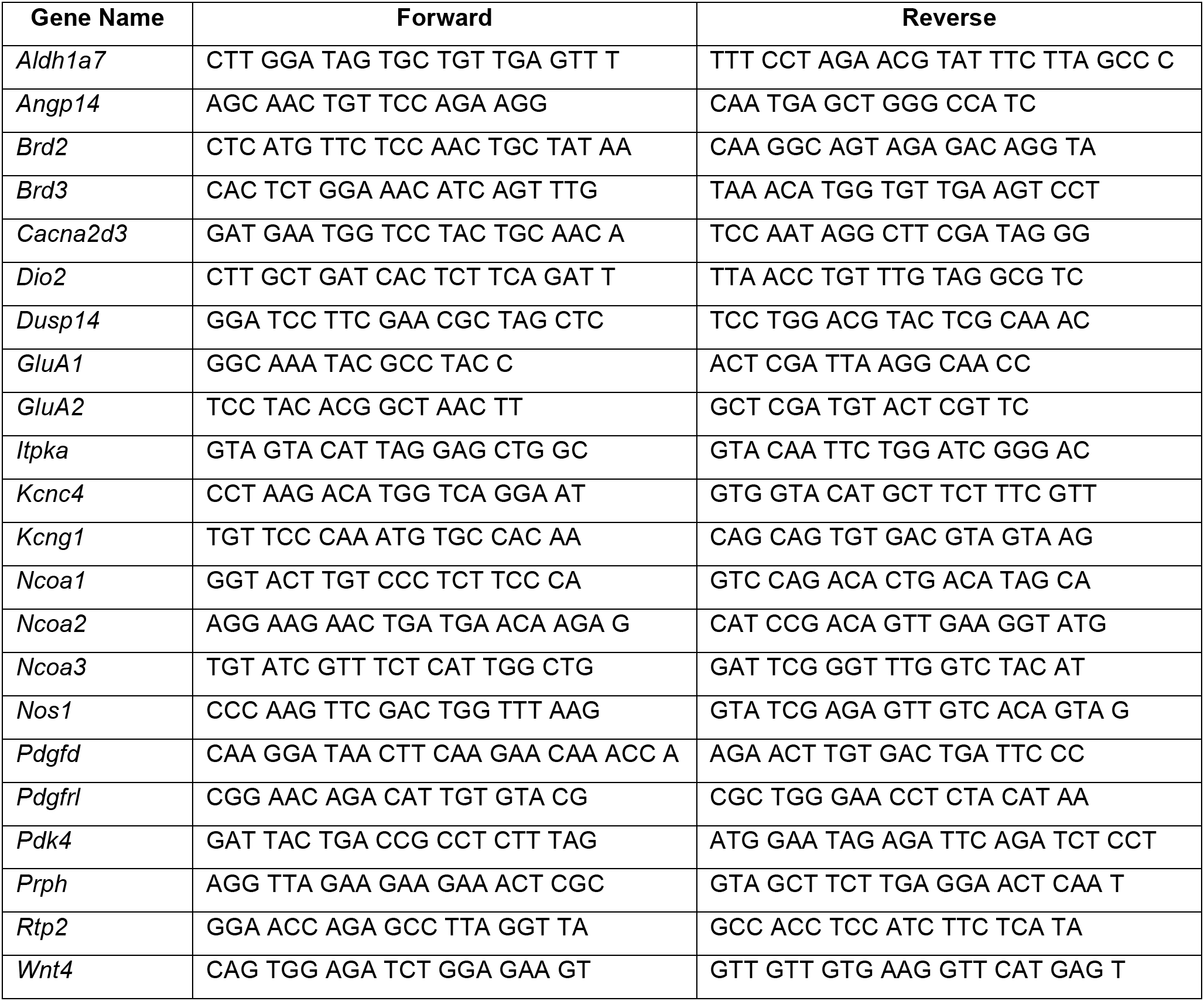
List of RT-qPCR primers

